# An Analytical Description for Action Potential Thresholds Defined by Concavity Changes

**DOI:** 10.64898/2026.04.21.719992

**Authors:** Marco Arieli Herrera-Valdez

## Abstract

A novel mathematical framework to define the threshold of action potentials in excitable cells is presented. Unlike previously applied methods that rely on approximations or specific fixed-point bifurcations, the approach focuses on the geometry of membrane potential trajectories. Specifically, the focus is on the concavity changes during the upstroke of an electrical pulse. These changes in concavity form a curve of inflection points that defines a region in phase space crossed by all the action potentials in the system, and containing no non-action potential trajectories. Such region is called the excitability region and its size can be measured, thus providing a measure for the excitability of a dynamical system, and a way to compare the excitability between systems representing different biological phenotypes and stimulus conditions. The work transforms the traditionally vague physiological concept of excitability into a rigorous analytical description applicable across continuous, single compartment models of electrical excitability.

## 1 Introduction

Electrically excitable cells are capable of producing all or none, pulse-shaped deflections in their transmembrane potential, *v* (in mV) (Aidley, 1998). Such pulses named *membrane action potentials (APs)* in the seminal work of Hodgkin and Huxley (1952), participate in different physiological functions that include rapid transmission of signals (Harper and Lawson, 1985), secretion of neurotransmitters and hormones (Dunant, 1994), and regulation of protein expression that supports cellular architecture and plasticity (Desai et al., 1999; Marrone and Petit, 2002). Examples of electrically excitable animal cells include neurons (Cole and Curtis, 1938; Hodgkin and Huxley, 1952), neuroglia (Verkhratsky et al., 2020), striated muscle fibers (Clausen et al., 1998; Stephenson, 1998), cardiomyocytes (Huang and Lei, 2023), pancreatic acinar cells (Petersen, 1982) and *β*-cells (Chay, 1987), and adrenal chromaffin cells (Biales et al., 1976). Of note, there are also electrically excitable cells in plants (Pickard, 1973; Wayne, 1993).

The all-or-none nature of APs (Fig. 1) suggests that electrical excitability is a threshold phenomenon (Fitz-Hugh, 1955; Karreman, 1951). Mathematical analysis of membrane potentials and electrical excitability has been performed in search of theoretical descriptions of thresholds (Fitz-Hugh, 1960; Noble and Stein, 1966). One outcome from that body of work is a classification for different types of threshold phenomena proposed by Fitz-Hugh. However, no specific equations describing thresholds for APs have been obtained and further work has only proposed approximations as solutions. Of particular interest, Platkiewicz and Brette (2010) derived an equation that describes an approximation for a voltage-threshold by taking into consideration the contribution of sodium currents to the upstroke velocity in action potentials. The impossibility of describing thresholds for voltage or current in models of neuronal membrane potential has also been discussed by other authors (Koch et al., 1995). From that work it has been argued that it is possible to determine a voltage-threshold separatrix curve in 2-dimensional models of membrane potential, but only if the underlying dynamical system has a saddle-node fixed point (Izhikevich, 2007); the separatrix curve is actually the unstable manifold of the saddle point. Another kind of boundary separating action potentials from other trajectories in phase space can be found in bistable systems having an attractor fixed point and an attractor limit cycle; in this case there is an repeller limit cycle surrounding the fixed point that can be thought of as a separatrix between action potentials (trajectories converging to the limit cycle) and non-action potentials (trajectories converging to the fixed point). However, the repeller cycle cannot be described analytically in biophysical models.

**Figure 1:**
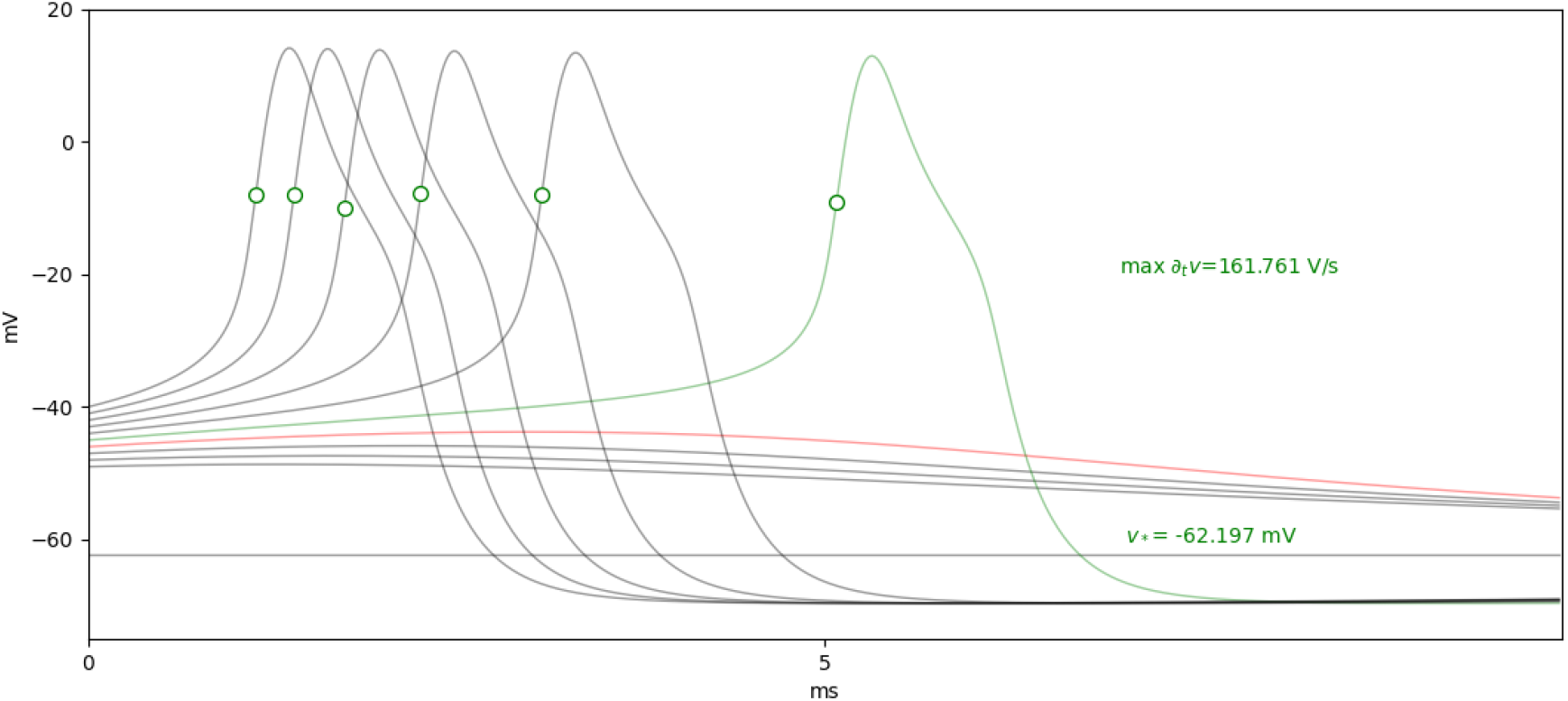
Point of no return from different initial conditions and in the absence of current stimulation (*a*_*F*_ = 0) assuming *w*(0) = *w*_∗_ with *v*(0) ∈ {−49, −41} in steps of 1 mV. The green circles mark the point at which the maximum rate of change with respect to time occurred for each trace. The green and red traces show the transitions between APs and non-APs (at a resolution of 1 mV). The green dots show the points at which the maximum ∂_*t*_*v* is reached during APs.

The discussion about thresholds can be clouded by the fact that models of membrane potential based on autonomous equations cannot be thought of as single dynamical systems, but instead, families of dynamical systems. The reason is that one autonomous dynamical system^1^ includes the trajectories that can be produced by fixing all the parameters in the equations. However, consider a model aims to reproduce results obtained from a current-clamp experiment such that the input current is changed from 0 to a non-zero value, and then back to 0, while the membrane potential is being recorded. The voltage trajectories in the model should then include trajectories like those produced in response to different amplitudes of current, which would require varying the current amplitude parameter, which means dealing with a family of dynamical systems (one for each value of the forcing parameter). Excitability has been classified in this case by essentially distinguishing qualitatively different types of transitions in the current-frequency (I-F) relationship between rest and repetitive firing as a function of the input current (Hodgkin, 1948). The main distinction between I-F relationships describes type-I excitability as a having a “continuous” I-F curve, whereas type-II excitability is related to a “discontinuous” I-F curve.

It is worth noticing that, from a mathematical stand point, the transitions between rest and repetitive firing occur at bifurcations in the underlying dynamical systems with the input current as a bifurcation parameter, always involving the emergence of an attractor limit cycle, which by definition has an intrinsic oscillation frequency that cannot be arbitrarily close to 0. In other words, there is no such thing as a continuous I-F curve. One additional issue is that the effect of increasing the stimulus amplitude within a large enough interval is to change the number or the types of fixed points, and their attractivity (Herrera-Valdez, 2012, 2018). It is therefore important to consider the thresholding problem in the different regimes defined by different stimulus amplitudes.

### What is excitability?

An electrically excitable dynamical system can be defined in simple terms as having a partition of its trajectories into two sets (Fitz-Hugh, 1955), those describing APs, and its complement (Fig. 1, red and green traces). Closer examination of the trajectories that are APs shows that the upstroke starts with a downward to upward change in concavity, and later changes back to downward as it passes through the point with the maximum rate of change with respect to time, before reaching the peak (Fig. 1, green trace, and Fig. 2). In both of the points just mentioned, 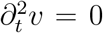. For a neuron, the local maximum for ∂_*t*_*v* may reach values beyond 100 mV/ms, which allows a practical distinction between trajectories that are APs and trajectories that are not (Fig. 1, red trace). As a consequence, trajectories that are not APs do not exhibit a large acceleration and a up-down change their time-curvature before they reach their peak. This suggests that the set of inflection points for the trajectories of the system may contain the information that is necessary to define a threshold, and also, such a set should allow partitioning the trajectories into two sets: those that are, and those that are not APs, thereby describing whether the system is excitable or not.

**Figure 2:**
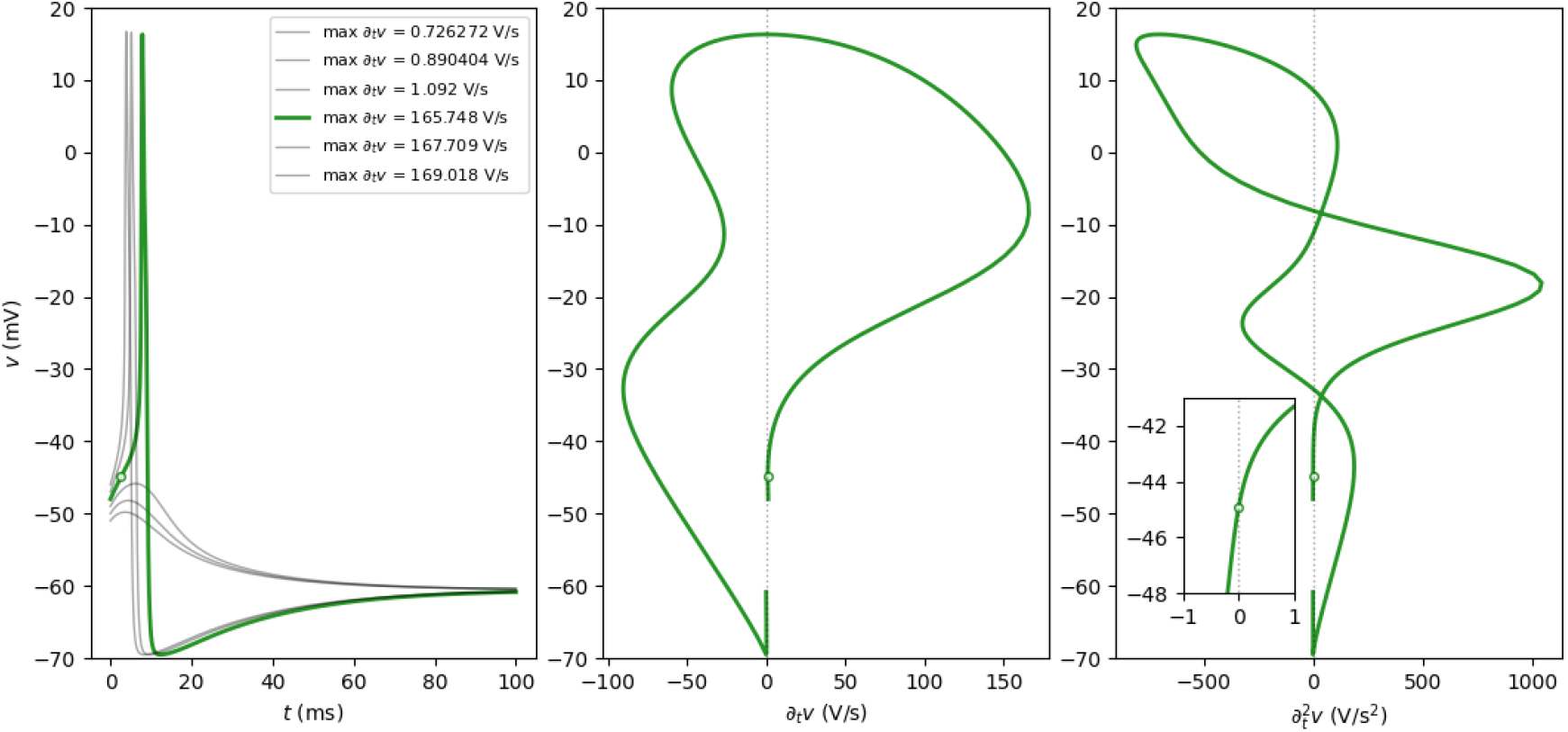
Move down to the section with the 2D model, and insert a figure here with the time-dependent dynamics for ∂_*t*_*v* and 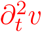. Rate of change and curvature profiles during an action potential. From left to right, (*t, v*), (∂_*t*_*v, v*), and, 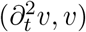. The green dot filled with white shows the threshold crossing, and the inset in the 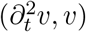 plane shows the curvature change at the time of threshold crossing. It was assumed that *a*_*D*_ = 19 and the rest of parameters as found in Table 2.

The curve describing the inflection points in phase space should be a manifold intersecting the upstrokes of the trajectories, marking points of no return for each trajectory. In phase planes, the AP thresholds can be thought of as curves intersecting the flow of trajectories on their way up. Trajectories that do not cross such points, do not continue increasing and return to rest.

To describe this phenomenon analytically, we consider first a model of neuronal membrane potential based on biophysical, and therefore continuous formulations. Since the second derivative with respect to time for *v* is a function of at least two variables in any such model, the thresholding will be studied in the phase space of the system. The initial focus will be to search for a “threshold set” assuming that all parameters of the models are fixed, with the stimulus amplitude equal to 0. In other words, we start by studying one dynamical system. Having done that, the description will be expanded to the whole family of dynamical systems included in the model. In particular, the analysis includes a description of how the thresholds given by inflection points during the upstroke change as the stimulus amplitude increases, as would be the case in a current-clamp experiment.

## 2 Membrane potential and curvature

Consider a general model of neuronal membrane dynamics. Written in general terms, such a model should include a variable *v* ∈ (*v*_*m*_, *v*_*M*_) (in mV) representing the transmembrane potential, a set of complementary variables represented by a vector 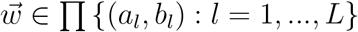 representing gating variables for ion-channels and possibly concentrations of molecules of interest (Best et al., 2007; Rasmusson et al., 1990; Wilson et al., 2004), and a vector 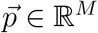 of parameters, with dynamics given by

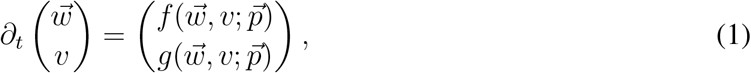

with functions *f* and *g* assumed to be smooth. If we assume further that *f* and *g* are derived from biophysical principles, *f* can be thought of as a non-linear function representing the difference between an input current *J*_*F*_ and the sum of voltage-dependent transmembrane currents, all normalized by the membrane capacitance^2^.

### Curvature changes and AP thresholds

From here one, thresholds are referred to as points that trajectories cross. If and when that happens, thresholds are points of “kinetic commitment”, during the upstroke which might be local minimae in ∂_*t*_*v* at the beginning of an AP (point of no return), or a local maximum of ∂_*t*_*v* before the peak that yields a landmark to detect the AP and its time.

In contrast, a separatrix is thought of here as a boundary in phase space that separates trajectories into two sets: APs and non-APs. Notice that the commitment of a trajectory to firing an AP occurs as the trajectory increases and becomes concave-up with respect to time (green traces in Fig. 1, and in Fig. 2). APs are concave down at first (near the resting potential), they cross the threshold when the cross the inflection point that changes the concavity upward (Fig. 2, see inset in the 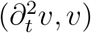 plane). In contrast, non-AP trajectories start concave-down with respect to time and do not change concavity as they return to the resting potential (Fig. 1, red trace).

Assume that vector of parameters 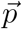 is fixed, and consider the resulting dynamical system with evolution rules given by equation (1). The curvature of *v* with respect to time in the resulting dynamical system is described by the set

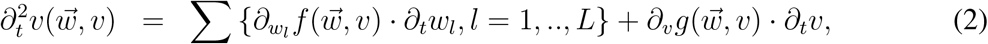

and the 0-level curve for 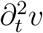,

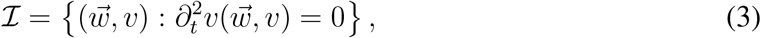

is a manifold in the 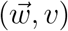-space that contains the inflection points of the different trajectories of the system.

#### Thresholds

Assume that 𝒯 is a trajectory in phase space, that describes an AP when projected to the (*t, v*)-axis. The AP peak is the point where 𝒯 crosses the *v*-nullcline 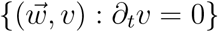 𝒯∩ℐ is the set of inflection points of the trajectory, and the AP threshold is the inflection point at which the minimum ∂_*t*_*v* occurs during the upstroke. That is, the point where 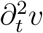 changes from negative to positive during the upstroke (Fig. 2).

If a trajectory describes more than one action potential, as it is the case for trajectories converging toward a limit cycle in these kinds of systems, then there should be more than one intersection of the upstroke of the trajectory with the set ℐ (one for each AP in the trajectory). A couple of questions that come up in anticipation of the analysis is how the sets containing the AP thresholds and the separatrix would change as the input current increases? How do these sets change if other parameters change? (say, if the number of K-channels in the membrane changes).

The discussion will focus on neuronal membrane potentials from this point on, but the results can be generalized to models of membrane potential for other excitable cell types.

### 2.1 Biophysical 2-dimensional description for membrane dynamics

The biophysical models of neuronal membrane potential *v* of the lowest dimension that can be constructed with continuous, autonomous equations are 2-dimensional. Such models typically assume that the membrane dynamics are generated by currents genrated by Na and K ions (Av-Ron et al., 1991; Fohlmeister et al., 1990; Rinzel, 1985). A second variable *w* replacing the vector 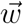 can be though of as the proportion of activated voltage-gated K-channels. These dynamics can be explicitly described by equations of the form

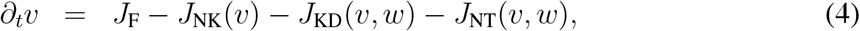

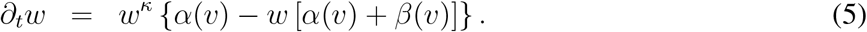

for *w* ∈ (0, 1) and *v* ∈ (*v*_K_, *v*_Na_), with *v*_K_ and *v*_Na_ representing the Nernst potentials for K^+^and Na^+^, respectively. To be able to compare APs in phase space with APs in the (*t, v*) plane, all the phase planes will be shown with the *w* in the horizontal axis and *v* in the vertical axes, respectively. That is, all the systems taken into consideration from here on will be defined within the domain 𝒟 = [0, 1] × [*v*_K_, *v*_Na_].

The terms *J*_NK_, *J*_KD_ and *J*_NT_ represent voltage-dependent transmembrane currents (all normalized by the membrane capacitance *C*_*m*_ (Herrera-Valdez, 2020), in units of A/F) generated by the Na-K ATPase, non-inactivating potassium (*a*.*k*.*a*. kalium) delayed rectifier channels, and transient sodium (*a*.*k*.*a*. natrium) channels, respectively (Herrera-Valdez, 2018). *J*_*F*_ (in A/F) represents a current stimulus amplitude, after normalization by *C*_*m*_. Note that *w* also represents the proportion of inactivated Na^+^channels, so that 1 − *w* represents the proportion of non-inactivated Na^+^channels. If *κ >* 0 and *v* is fixed, the term *w*^*κ*^ yields solutions for *w* with logistic shape, as shown in experimentally recorded currents (for instance, see Fig. 3 in the classical paper by Hodgkin and Huxley (1952) and in Tsunoda and Salkoff (1995)).

**Figure 3:**
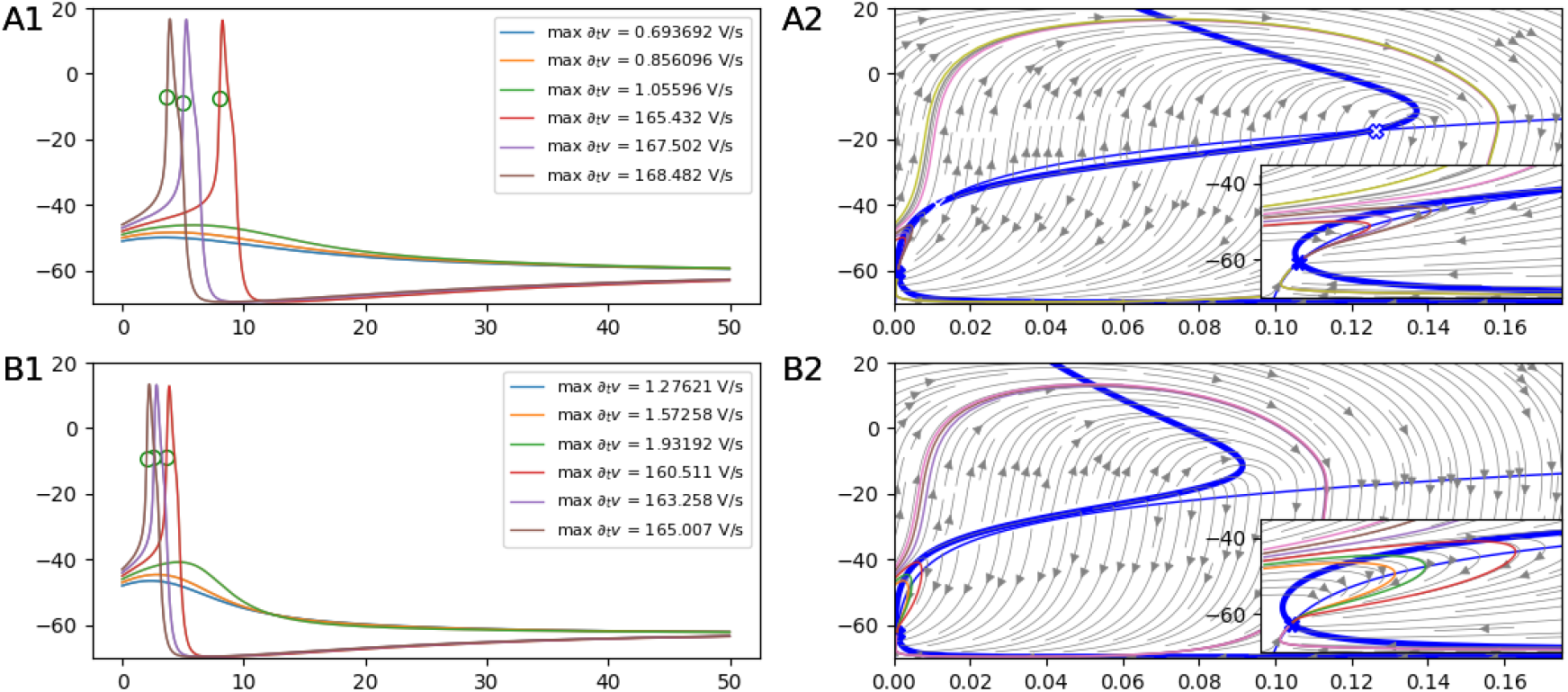
Two phase planes illustrating the two topologically different configurations of systems from equations (4)-(11) in the absence of repetitive spiking, assuming *J*_*F*_ = 0. The thick and thin blue lines show the *v*- and *w*-nullclines, respectively. The attractor nodes are marked as blue circles, the saddle-node with a blue cross, and the blue circle with a white cross is a repulsive node. (A1-A2) Three fixed points (*a*_KD_ = 19). (B1-B2) One fixed point (*a*_KD_ = 28). The fixed points that correspond to the resting potential are marked as blue squares. Insets in A2 and B2 show the dynamics near the attractor point. In the configuration with 3 fixed points (A1-A2), there is also a saddle-node marked with an “x”, and a repeller focus point marked as an empty circle.

Explicitly, the currents can be written in general terms as

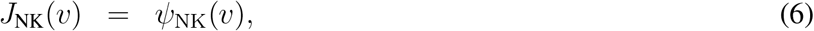

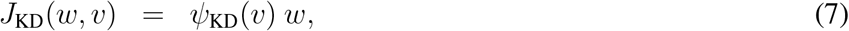

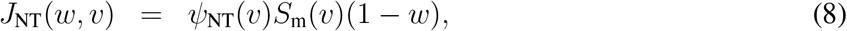

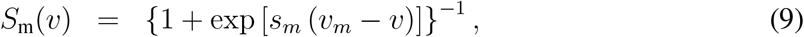

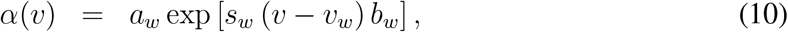

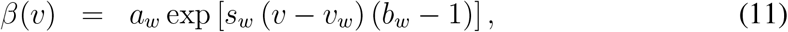

with the amplitude term *a*_*w*_ in units of 1/ms. The terms *ψ*_*l*_, *l* ∈ {NK, KD, NT} represent the driving force for the currents. Using the thermodynamical model for transmembrane transport ^3^ (Herrera-Valdez, 2018), the transmembrane currents currents can be written

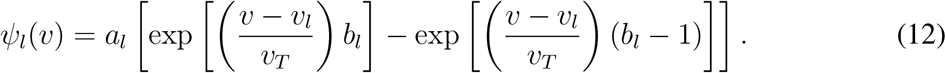

with *a*_*l*_ = *A*_*l*_*/C*_*m*_ (pA/pF, *A*_*L*_ in pA, *C*_*m*_ in pF), for *l* ∈ {NK, KD, NT}. The thermodynamical model can be simplified sacrificing the first principles approach for the sake of mathematical tractability. The conductance-based model used by Hodgkin and Huxley (1952) is a linear approximation for *ψ*_*l*_ around the reversal potential *v*_*l*_ (Herrera-Valdez, 2018) that yields

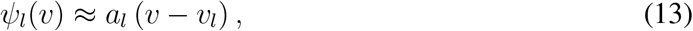

with *a*_*l*_ = *A*_*l*_*/*(*C*_*m*_*v*_T_) (1/ms). The example model used to illustrate the calculations will be conductance-based, but the results shown here hold for the thermodynamic model too.

From a dynamical systems point of view, *v* can be regarded as a fast and self-amplifying variable that positively feeds into a slower variable *w* that, in turn, negatively feeds back into *v*. The slower dynamics of *w* allow some the trajectories in *v* to increase before causing it to decrease.

#### 2.1.1 Geometry underlying the 2D dynamics

To be able to compare trajectories in the (*t, v*) plane with those in the phase plane, all phase planes will be shown with the *w*-coordinate in the horizontal axis and the *v*-coordinate vertically (Fig. 1).

Notice that the equations (4)-(11) can be rewritten as

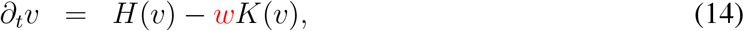

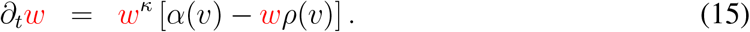

with

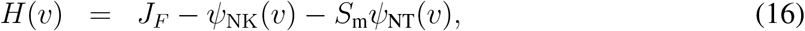

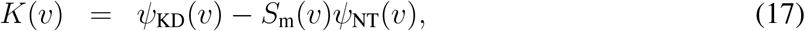

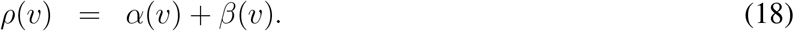

From there, it is possible to obtain analytical expressions for the nullclines, curvature, and determine the stability and type of the fixed points, also analytically.

##### Nullclines and fixed points

Using equations (14)-(15) the nullclines are the sets

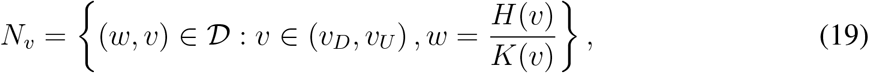

and

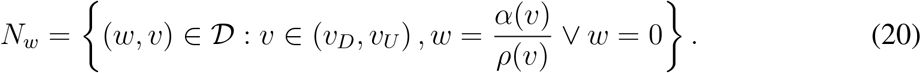

The phase plane is divided by the nullclines into four regions in which the signs of ∂_*t*_*v* and ∂_*t*_*w* alternate, as described in Table 1. The parameters must be such that the *v*-nullcline does not cross the line {(*w, v*) : *w* = 0}.

**Table 1.** Regions of the phase plane defined by the nullclines. The discussion of this paper about thresholds focuses on *R*_1_, where all AP upstrokes are found.

| Region | Description | $\partial_t v$ | $\partial_t w$ | AP segments contained |
| --- | --- | --- | --- | --- |
| $R_1$ | $\left\{ (w, v) : 0 < w < \min \left( \frac{H(v)}{K(v)}, \frac{\alpha(v)}{\rho(v)} \right) \right\}$ | + | + | Upstroke, return to resting potential |
| $R_2$ | $\left\{ (w, v) : \frac{H(v)}{K(v)} < w < \frac{\alpha(v)}{\rho(v)} \right\}$ | - | + | Initial Downstroke |
| $R_3$ | $\left\{ (w, v) : \max \left( \frac{H(v)}{K(v)}, \frac{\alpha(v)}{\rho(v)} \right) < w < 1 \right\}$ | - | - | Afterhyperpolarization |
| $R_4$ | $\left\{ (w, v) : \frac{H(v)}{K(v)} > w > \frac{\alpha(v)}{\rho(v)} \right\}$ | + | - | Initial repolarization |

Note that $R_1$ , the region where both variables increase, is where all the upstrokes for APs are found. Most of the discussion will focus on this region. The set of *fixed* points of the system is the intersections of the two nullclines. That is, the set $N_v \cap N_w$ .

**Table 2.** Parameters for the model. The reversal potential for the Na-K ATPase current is given by *v*_NK_ = 3*v*_Na_ − 2*v*_K_ + *v*_ATP_. All the rate amplitudes for the currents are maximal membrane conductances normalized by the product of *v*_T_ and the membrane capacitance (Herrera-Valdez, 2018).

| Parameter | Value/Range | Units | Meaning |
| --- | --- | --- | --- |
| **Constants** |  |  |  |
| $k_B$ | $1.380649 \times 10^{-23}$ | Joules/Kelvin | Boltzmann's gas constant. |
| $q_e$ | $1.60217663 \times 10^{-19}$ | Coulombs | Elementary charge. |
| $T_C$ | 36.5 | ° Celcius | Temperature |
| $T$ | 309.65 | Kelvin | Absolute temperature (Kelvin, $T = 273.15 + T_C$). |
| **Voltages** |  |  |  |
| $v_{\text{T}} = k_B T / q$ | 26.7 | mV | Thermal potential. |
| $v_{\text{K}}$ | -90 | mV | Nernst potential for $\text{K}^+$ |
| $v_{\text{Na}}$ | 60 | mV | Nernst potential for $\text{Na}^+$ |
| $v_{\text{ATP}}$ | -430 | mV | ATP hydrolysis potential |
| $v_w$ | 0 | mV | Half-activation potential for KD channels |
| $v_m$ | -20 | mV | Rectification bias for NaT channels |
| **Gain factors and biases** |  |  |  |
| $b_w$ | 0.65 | – | Activation bias for KD channels |
| $s_w$ | $3.5/v_{\text{T}}$ | 1/mV | Gain for KD channel activation, normalized by $v_{\text{T}}$ |
| $s_m$ | $5.5/v_{\text{T}}$ | 1/mV | Gain for NaT channel activation |
| **Rates** |  |  |  |
| $a_{\text{NK}}$ | 0.3 | 1/ms | Rate (amplitude) for the Na-K ATPase current |
| $a_{\text{KD}}$ | 19, 28 | 1/ms | Rate (amplitude) for the current mediated by KD channels |
| $a_{\text{NT}}$ | 3 | 1/ms | Rate (amplitude) for the current mediated by NaT channels |
| $a_w$ | 0.1 | 1/ms | Rate for the activation of KD channels |

Note that *R*_1_, the region where both variables increase, is where all the upstrokes for APs are found. Most of the discussion will focus on this region. The set of *fixed* points of the system is the intersections of the two nullclines. That is, the set *N*_*v*_ ∩ *N*_*w*_.

##### A note on bifurcation structures

If the system does not already exhibit repetitive spiking, then it has three configurations, with three, two, and one fixed point respectively. For small enough *J*_*F*_ (before transitioning into repetitive spiking), all three of these conditions are such that the fixed point with the smallest *v*-value is an attractor, and its *v* value corresponds to the resting potential *v*_*r*_. The configuration with two fixed points is very unlikely to be observed, both in experiments and numerically. It occurs as a transition between three and one fixed points. One way to observe the transition is to start with three fixed points, and to increase the stimulus amplitude *J*_*F*_, which will eventually lead to a saddle-node bifurcation, in which two fixed points will be lost as they collide before disappearing. Another way to observe the transition, now assuming *J*_*F*_ = 0, is to increase *a*_KD_, the amplitude of the K^+^current, which is equivalent to think of a membrane with more K^+^channels. The transition between these two configurations can also be achieved by changing any parameter that affects the slope of *N*_*w*_, or the slope of the middle branch of *N*_*v*_. Two such parameters are *s*_*w*_ and *v*_*w*_.

It should be noted, however, that in both configurations, the sequence of fixed point bifurcations with respect to the parameter *J*_*F*_ includes a subsequence switching attractor nodes, to attractor focus, repeller focus, attractor focus, and attractor node, at least. Limit cycle attractors emerge through fixed point bifurcations (e.g. saddle-node on invariant circle, or supercritical Andronov-Hopf) or through limit-cycle bifurcations (e.g. saddle-node off invariant circle, or fold) that result in bistability with one attractor focus point and an attractor limit cycle (Herrera-Valdez, 2012; Herrera-Valdez et al., 2013).

##### Inflections during the upstroke as thresholds

The inflection points can be calculated from equations (2), (14), and (15). The idea is to take into account that *v* and *w* are functions of *t* and use the chain rule for differentiation. Explicitly,

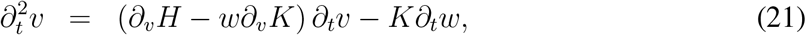

which set equal to zero yields the set

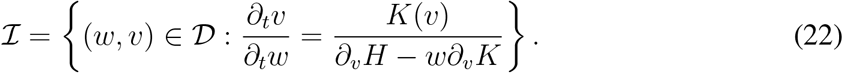

The set of points that we will be regarding as AP thresholds should be part of the AP upstroke and at such points there should be changes in the concavity of the *v*(*t*) curve, from positive to negative on its way to the AP peak. In other words, to find the upstrokes we need to find a subset of ℐ ∪ *R*_1_.

##### Geometry of action potentials and excitability

One of the main goals of this work is to define excitability from a geometrical stand point. If excitability is the capability of producing APs, then the definition would be incomplete until an AP is defined first.

So, if a trajectory in the phase plane is such that (i) it starts in region *R*_1_ and crosses ℐ before reaching the maximum point for *v* at *N*_*v*_; (ii) and continues through regions *R*_2_, *R*_3_, and *R*_4_, in that order, then such a trajectories will be regarded as an AP. As a consequence, the set of orbits in an excitable system can be partitioned into a set that only contains APs, as defined above, and its complement. In particular, bistable systems with one attractor fixed point and one attractor limit cycle are therefore excitable. In contrast, systems that already contain a unique limit cycle attractor can be referred to as already “excited”, instead of excitable.

#### 2.1.2 An example model

To illustrate the behaviour of the inflection points, we will calculate the curvatures for the model in equations (4)-(11), assuming *a*_*NP*_ = 0 = *κ*, assuming conductance-based approximations for current, with parameters as in Table 2. The behaviour of the inflection points in the absence of stimulation (single dynamical system with *J*_*F*_ = 0, Fig. 3) is studied first, taking into consideration the two configurations of the system with three fixed point (*a*_KD_ = 19) and one fixed points (*a*_KD_ = 28). The behaviour of the inflection points for other dynamical systems of the model (different values of *J*_*F*_) is considered after.

##### Model characteristics and mathematical constrains for parameters

The APs in the model dynamics should be such that maximum 100 ≤ max {∂_*t*_*v*} ≤ 300 V/s, lasting between 1 and 3 milliseconds, with amplitudes between 80 and 100 mV (Fig. 1), with a rheobase between 25 and 100 pA.

Note that the function *K*(*v*) is positive for all *v* ∈ (*v*_K_, *v*_Na_) because the driving forces for K^+^and Na^+^are respectively positive and negative within the interval (*v*_K_, *v*_Na_). Also, *ψ*_KD_(*v*) is positive, while *ψ*_NP_(*v*) and *ψ*_NT_(*v*) are negative for all *v* ∈ (*v*_K_, *v*_Na_). However, *ψ*_NK_ can be both positive and negative for *v* ∈ (*v*_K_, *v*_Na_), as it changes sign at *v* = *v*_NK_. Taking the above observations into consideration, the amplitude parameters for the currents should be such that the *v*-Nullcline does not cross the *v*-axis around *v*_NK_, the reversal potential for *ψ*_NK_. In more general terms, the parameters of the model should be such that the function *H*(*v*)*/K*(*v*) that defines the *v*-nullcline (equation (19)) yields values for *w* in the interval [0, 1].

The calculations for curvatures and inflection points will be performed using the model in its general form using equation 21. We will use this simplified model to illustrate the thresholding calculations with the example model. See Table 2 for more details.

#### 2.1.3 Refinement of the conjectures for the 2D-geometry

Assume that the parameters are fixed and such that the resulting system does not have a limit-cycle attractor (no repetitive firing). Then it should be the case that: (i) For APs starting near the resting potential, the AP threshold (along a single trajectory 𝒯) is the inflection point in *R*_1_ where the concavity changes from downward to upward 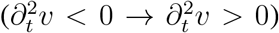. (ii) For all APs, there should be an inflection point before the peak where the concavity changes from up to down 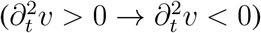. (iii) As a consequence of (i) and (ii), there is a subset *E* ⊂ *R*_1_ containing all the orbit segments of trajectories that are APs, and containing no non-AP orbits. The vertical boundaries of such a set include the inflection points mentioned in (i) and (ii). The existence of such a set, and its size within the domain 𝒟 = [0, 1] × [*v*_K_, *v*_Na_], can then be used to decide if a system is excitable, and measuring its size should provide a way to compare excitability between systems.

## 3 Results

Assume that all the parameters in equations (14)-(18) are fixed with *J*_*F*_ = 0 (see Table 2). In this case the system is excitable, which means that some of its trajectories are action potentials, with one attractor fixed point whose *v*-value represents the resting membrane potential. Recall that the system is likely to be in one of two possible configurations (i.e. two types of excitable systems) corresponding to three, or one fixed point respectively (Fig. 3). Note that the *w*-values in which the trajectories for the system with three fixed points occur (Fig. 3A2) is larger than the corresponding *w*-values for the system with one fixed point (Fig. 3B2). In other words, the level of Na-channel inactivation, or analogously, the level of K-channel activation for the 3-fixed point configuration is larger in comparison to the Na-inactivation in the 1-fixed point regime. This can also be observed in the duration of the upstrokes of the action potentials for the two systems (Fig. 3A1,B1), all of which were calculated starting from the same region.

### 3.1 Curvature changes and thresholds for APs

From the calculations of curvature in equation (21), the 0-level curve of 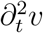 (Fig 4, green lines) is the set of inflection points given by

**Figure 4:**
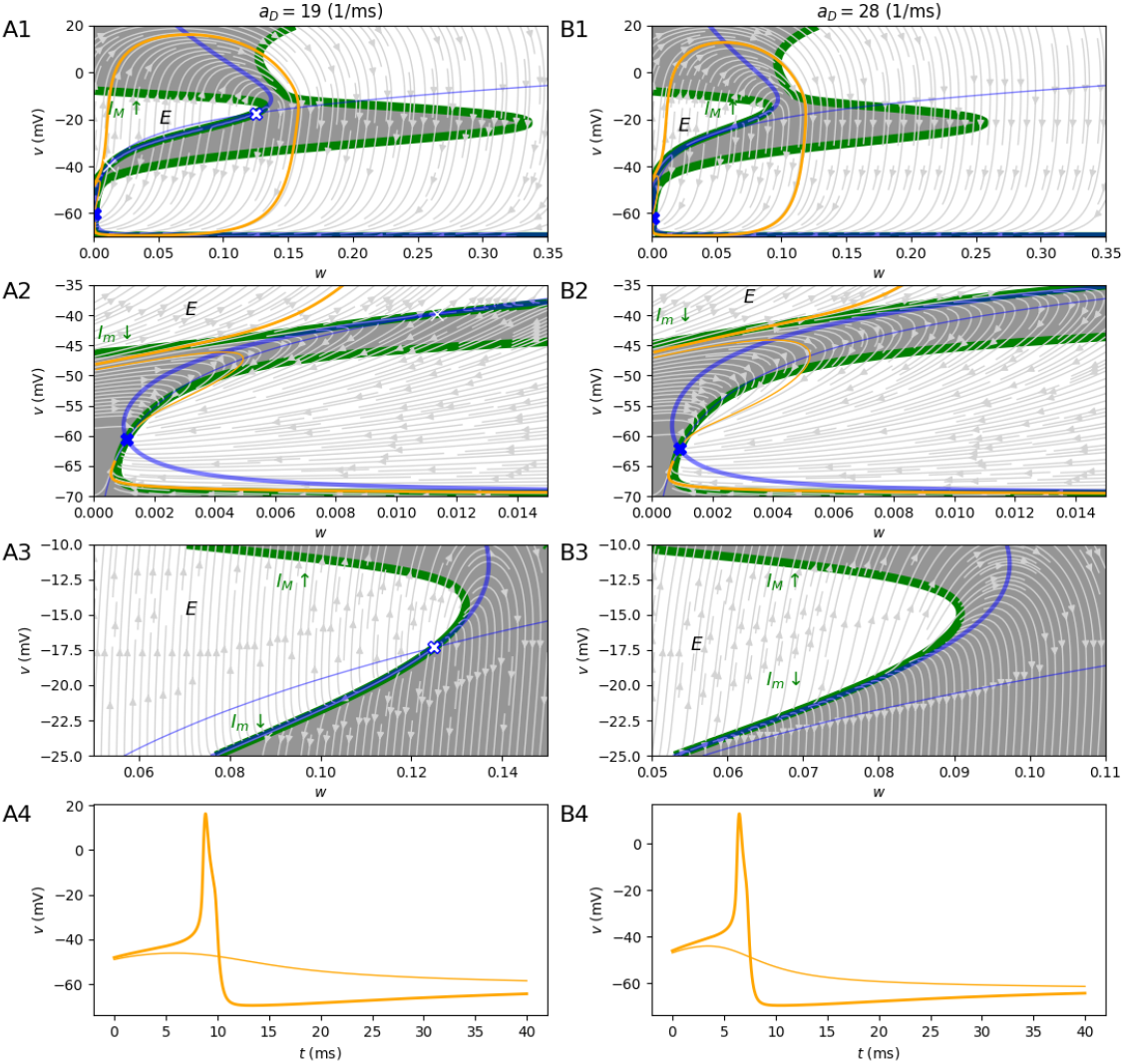
Curvature in the phase plane and APs that change curvature twice, for *a*_*D*_ ∈ *{*19, 30}. The green lines represent the inflection point curve ℐ. The white and gray areas are zones where 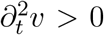 and 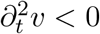, respectively. The thick and thin blue lines show the *v*- and *w*-nullclines, respectively. APs starting with 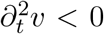 are shown by orange lines. A1-A4. System with three fixed points (*a*_KD_ = 19). B1-B4. System with 1 fixed point (*a*_KD_ = 28). A2 and B2 show the regions around the resting potentials and the division between non-APs and APs. A3 and B3 show the apex of ℐ_*θ*_ and non-APs (trajectories that do not cross the segment *I*_*m*_). Note that the region *E* is bounded in part by *N*_*w*_ when there are 3 fixed points. Notice APs always cross *I*_*M*_ and can be found for both configurations of the system. A4 and B4 show the trajectories in the phase plane starting below *I*_*m*_, with a downward curvature. Include maximum rates of depolarization in the plots with APs.

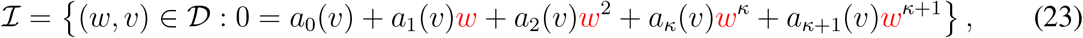

ith

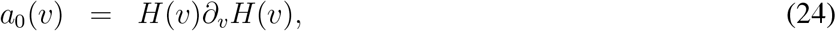

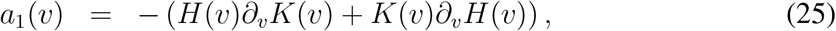

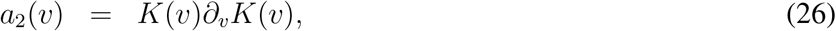

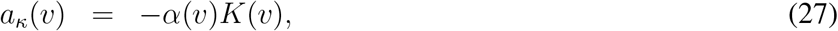

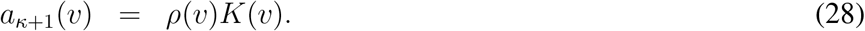

#### Special case: quadratic solutions for *κ* = 0

Note that the curve ℐ can be calculated explicitly as a function of *v* when *κ* = 0. In that case the coefficients transform into

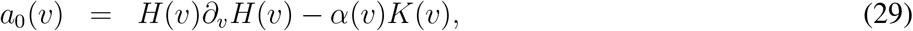

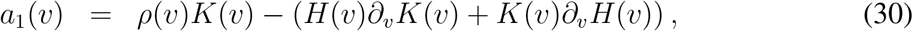

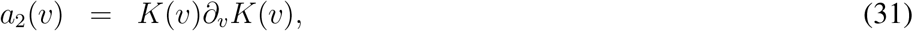

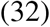

and the solutions in that case would be

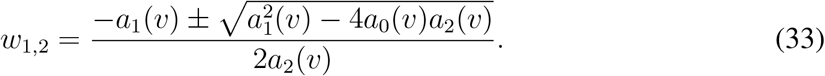

#### 3.1.1 Curvature for *J*_*F*_ = 0

the curve ℐ of inflection points has two connected components in 𝒟 for *J*_*F*_ = 0, for the two configurations of the system (Fig. 4, green lines). The right-most component is crossed by all the segments of trajectories that either go down (∂_*t*_*v <* 0), or go toward the attractor point from below (∂_*t*_*v >* 0) (Fig. 4A2-B2).

##### The threshold set

The left-most component of ℐ separates *R*_1_ vertically into three regions: two sets where 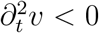 (Fig. 4, gray patches within *R*_1_) flanking a set where 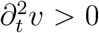 (Fig. 4, white patches). The subset ℐ_*θ*_ = ℐ ∩ *R*_1_ of the inflection curve ℐ separates these regions and intersects segments of APs. Note ℐ_*θ*_ is the union of an upper segment *I*_*M*_ and a lower segment *I*_*m*_ (Fig. 4A1,B1). The upper segment separates upward from downward curvatures with respect to time, intersects trajectories as they reach their maximum rate of change in the upper portion of ℐ_*θ*_ and change to a downward curvature on their way to the peak as they cross *I*_*M*_. In contrast, *I*_*m*_ is crossed by upstrokes that start near the resting potential, with an initial concave-down segment that becomes concave up with respect to time (Fig. 4A2,B2, and C). Those trajectories reach their minimum 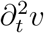 as they cross *I*_*m*_ (Fig. 4A2,B2).

Notice that not all trajectories that start in *R*_1_ are APs (Fig. 4-A2, B2, C). Some of the trajectories starting below *I*_*m*_ never cross *I*_*m*_, which means they never switch curvature from downward to upward as a function of time. Those trajectories reach *N*_*v*_, their local maximum without crossing *I*_*m*_. These trajectories may increase initially, but they eventually turn down and continue toward the attractor point.

Note that not all AP trajectories start concave down (see trajectories starting above *I*_*m*_ near the *v*-axis). However, all APs cross the region ℐ_*θ*_ = *I*_*m*_ ∪ *I*_*M*_, and all AP trajectories cross *I*_*M*_. Therefore, the set of trajectories (excluding those starting at FPs) for *J*_*F*_ = 0 can be partitioned into two sets containing APs and non-APs, respectively. This happens for both configurations of the system, which means that both systems are excitable for *J*_*F*_ = 0.

Notably, the threshold manifold *I*_*m*_ is lower for *a*_*D*_ = 19 in comparison to *a*_*D*_ = 28, in agreement with the idea that less K^+^channels should lower the AP threshold.

##### Is there a separatrix?

All the non-AP trajectories that start in *R*_1_, start below *I*_*m*_, which means they start concave-down, increase in both *w* and *v* and reach their maximum in *v* (intersect the *v*-nullcline) on their way toward the resting point.

In the configuration for 3 fixed points, the non-AP trajectory starting at *R*_1_ with the maximum *v* reaches *N*_*v*_ before reaching the (middle) saddle-node fixed point. That is, the *v*-peaks of non-APs starting at *R*_1_ are bounded by the *v*-coordinate of the saddle fixed point. The reason is that the *w*-nullcline becomes the right-most bound for *R*_1_ at that point, and it is below *I*_*m*_. Therefore, for the 3-fixed point configuration the set of non-APs that start in *R*_1_ is bound on the right by the stable manifold of the saddle-node fixed point, as already found by other researchers (Izhikevich, 2007), (Fig. 4A3).

In contrast, for the configuration with 1 fixed point, the non-AP trajectory starting in *R*_1_ with the highest peak stays below *I*_*m*_. The highest of those non-AP peaks occurs near the right tip of ℐ_*θ*_, close to where *I*_*m*_ meets *I*_*M*_ (Fig. 4B3).

In both configurations, the set of non-APs starting at *R*_1_ is formed by trajectories that do not cross *I*_*m*_ from below, whereas APs are trajectories that do cross ℐ_*θ*_ at least once. So the boundary of the set of non-APs starting at *R*_1_ can be thought of as a separating set that renders the system as excitable (Fig. 4A2).

Note the separatrix is a boundary between trajectories, and therefore, it is not crossed by trajectories, unlike ℐ_*θ*_.

##### The AP threshold is raised by K-channel activation (alternatively, Na-channel inactivation)

The shape of *I*_*m*_ (Fig. 4 in both configurations of the system is such that higher values of *v* are required for higher values of *w*. That is, *I*_*m*_ can be thought of as an increasing function of *w*, which means that the AP threshold increases with activation of *K*-channels.

An alternative interpretation could be drawn from this particular model, and that is that Na-inactivation yields higher thresholds, in agreement with the intuition that the upstroke occurs when Na-channels activate.

##### K-channel activation decreases the maximum rate of depolarization threshold

The max ∂_*t*_*v* threshold segment *I*_*M*_ in ℐ_*θ*_ requires larger values of *v* for smaller values of *w* (Fig. 4). As a consequence, K-channel activation yields smaller rates of depolarization during an action potential. As before, this result could be interpreted as Na-channel inactivation yielding higher thresholds (higher values of *w* correspond to smaller values of *v* in *I*_*M*_).

### 3.2 Excitability in a single dynamical system

Consider the set

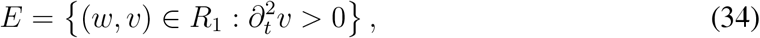

which is bounded by ℐ_*θ*_ (Fig. 4A1,B1).

Note that all of the AP upstrokes of the system pass through *E* before reaching their maximum ∂_*t*_*v* on their way to their peak. That is, all the trajectories in *E* are APs, and there is no AP that does not cross *E*. For this reason *E* will be called from here on, the *excitability set*, and a dynamical system with a region like *E* will be regarded as excitable from here on. By extension, a model composed of a family of dynamical systems can be called excitable if it contains excitable dynamical systems.

### 3.3 A measure of excitability

Can we say whether one of two of more systems is more excitable using the definition given above? In principle, it would be possible to compare excitabilities for dynamical systems if there was a way to measure their excitabilities.

To measure the size of the region *E* within the domain 𝒟, let

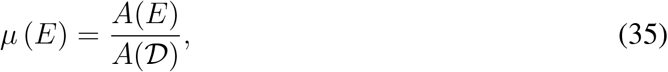

where *A*(·) represents the area. Note that *µ* can actually be regarded as a measure on 𝒟 that yields a number between 0 and 1, similar to a probability measure. Note that *A*(𝒟) = *v*_Na_ − *v*_K_.

For computational purposes, such a measure could be approximated by setting up a grid on 𝒟 with a fine mesh, count the number of points in *E*, and divide by the number of points in 𝒟. The process can be repeated by increasing the number of points in the mesh for 𝒟, and stopped when the difference between two values of *µ*(*E*) is less than some given tolerance.

#### 3.3.1 Comparing excitability across electrophysiological phenotypes

How can we use the measure *µ* to compare dynamical systems with different parameters? (*e*.*g*. model cells representing different electrophysiological phenotypes) For instance, is it possible to decide with precision whether neurons become more or less excitable by expressing more or less potassium channels, for the same level of external stimulation?

In principle, one condition should be that *A*(𝒟) is the same for all the systems under comparison. In general terms, consider two excitable dynamical systems *φ*_*C*_ and *φ*_*D*_ respectively corresponding to two sets of parameters *C* and *D* and regions of excitability *E*_*C*_ and *E*_*D*_ in the same domain 𝒟. Then we will say the system *φ*_*C*_ is less excitable than *φ*_*D*_ if *µ* (*E*_*C*_) *< µ* (*E*_*D*_).

For instance, in the examples shown above (Fig. 4), let *E*_19_ and *E*_28_ represent the excitability sets for the two systems with *a*_*D*_ ∈ {19, 28} and *J*_*F*_ = 0.

In the absence of other changes, *µ*(*E*_19_) *> µ*(*E*_28_), so the neuron with less K^+^channels (*a*_*D*_ = 19) is more excitable in comparison with the model neuron with more K^+^channels (*a*_*D*_ = 28).

#### 3.3.2 How does the curve ℐ_*θ*_ change by increasing *J*_*F*_ ?

For illustration, the evolution of the inflection curve ℐ_*θ*_ can be studied using different values of *J*_*F*_ (Fig. 5).

**Figure 5:**
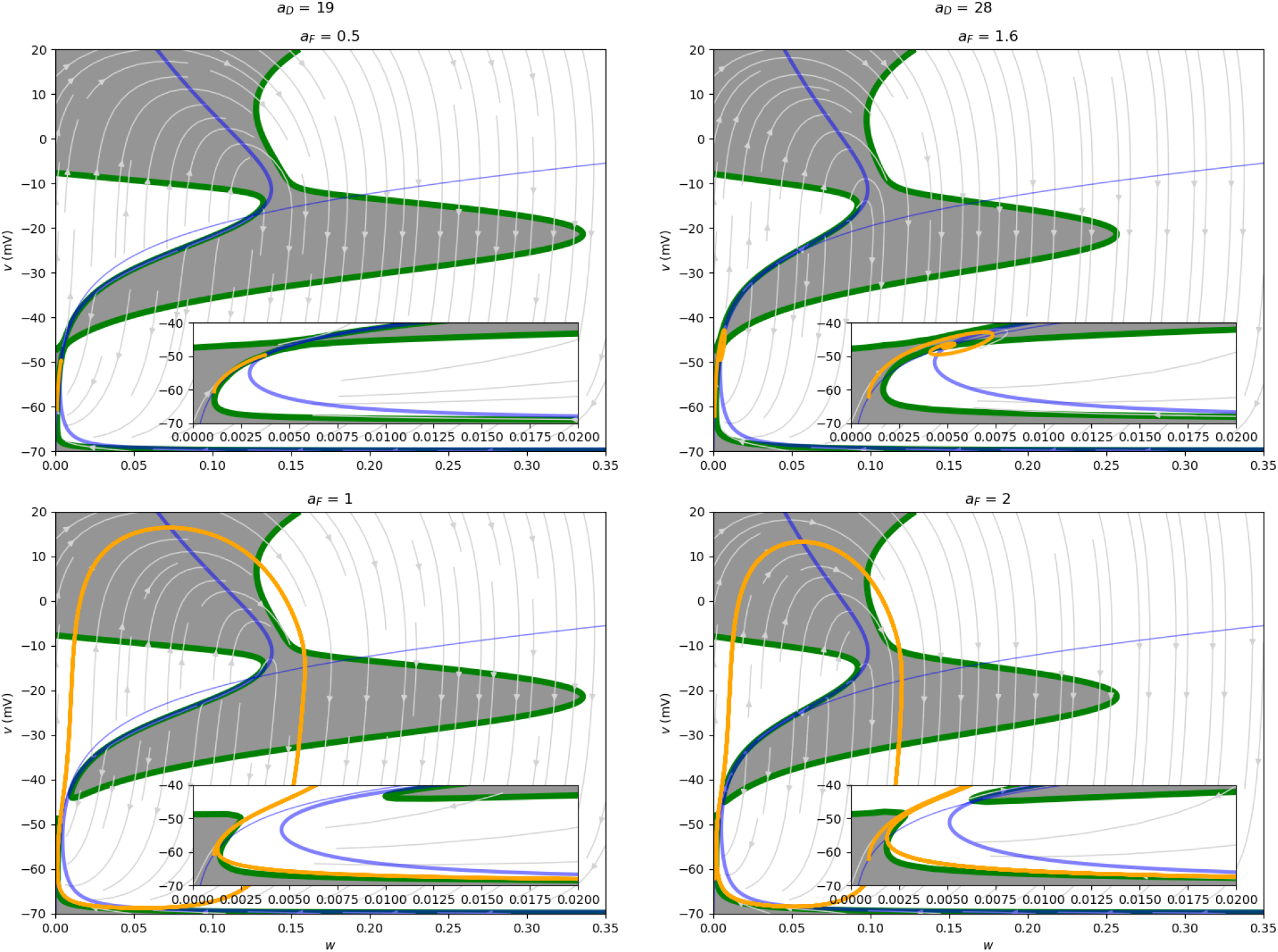
Progression of inflection point curves for the two configurations with 3 (left) and 1 fixed point (right). The orange curve shows one trajectory from (*w*_0_, *v*_0_) = ().

In the absence of other changes, in the two configurations of the system the curve of inflection points ℐ changes for increasing values of *J*_*F*_. For small enough values of *J*_*F*_, ℐ has two connected components, one of which is within the region *R*_1_. However, as *J*_*F*_ increases and the system stops being excitable and becomes excited (with a unique attractor limit cycle), the two components of ℐ change, and region where 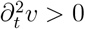 in *R*_1_ becomes fully connected with the rest of the phase plane (Fig. 5, insets).

It is also possible to compare the sizes of the excitability regions across systems. Let *φ*_*a,b*_ represent the dynamical system with values *a*_KD_ = *a* and *J*_*F*_ = *b*, with all the other parameters identical. Then from Fig. 5 it is possible to see that *µ* (*E*_19,0.5_) *> µ* (*E*_28,1.6_). So a neuron with more K^+^channels receiving a stimulus of amplitude *J*_*F*_ = 1.6 pA/pF may still be less excitable than another with less K^+^channels (*a*_*D*_ = 19) receiving a stimulus with amplitude *J*_*F*_ = 0.5 pA/pF.

## 4 Discussion

The results above show that there are thresholds for APs and that it is possible to write analytical expressions for such thresholds directly from continuous models underlying the dynamics, and relate them to biophysical parameters. The thresholds can be found by searching for inflection points in the AP upstroke. Explicitly, this is done by calculating the second time-dependent derivative of the membrane potential with respect to time, 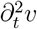, and finding its zero-level manifold ℐ. The intersection ℐ with the set containing AP upstrokes yields a manifold ℐ_*θ*_ with two segments *I*_*m*_ and *I*_*M*_ that contain points that show two different thresholds for APs. One of these segments, the threshold manifold *I*_*m*_ contains points of no return at which AP trajectories change their concavity from downward to upward, effectively starting the acceleration of the AP upstroke.

### Distinguishing APs and non-Aps

The segment, *I*_*M*_ contains the points at which all the APs in a system reach their maximum rate of depolarization. In other words, *I*_*M*_ contains all the points that represent the “all” part of the AP.

The region immediately below *I*_*m*_ within *R*_1_ in the phase plane only contains segments of trajectories in which *v*(*t*) is concave-down and never crosses *I*_*m*_. The closure of the set of trajectories in *R*_1_ that start concave down and never cross *I*_*m*_ is the set of non-APs. If this set contains at least one multi-point trajectory, then we can say there is a non-trivial partition of *R*_1_ into subsets containing segments of APs and non-APs.

One important property of the threshold curve ℐ_*θ*_ is that its dependence on states (*w, v*) agrees with the intuitive idea that K-activation (and Na-inactivation) increases the AP-threshold, but decreases the maximum rate of depolarization. However, Na-channel inactivation is not necessary to increase the AP threshold, as can be observed in a minimal “Up-Down” version of the model presented here, with only two non-inactivating voltage-gated currents, one pushing *v* upward (Na-current, *J*_Na_(*v*) = *a*_NP_*S*_NP_(*v*)*ψ*_NP_(*v*)), and the other pushing *v* downward (K-current) that yields AP threshold curves with similar shapes as the *I*_*m*_ and *I*_*M*_ curves presented previously.

### The excitability region can be used to measure excitability

The threshold manifold ℐ_*θ*_ defines a region *E* that contains trajectory segments belonging only to APs, and contains them all, so an excitable dynamical system is defined as one in which the phase space contains a region *E* as described above, and 𝒟 \ *E* is not discrete (contains at least one trajectory formed by more than one point).

Once the definition of excitability based on this analytical threshold set is provided, a measure of excitability is constructed, thus providing means to quantify excitability in single dynamical systems, and enabling the possibility of deciding whether one system is more excitable than another.

Can the measure of excitability proposed here be used to assess changes in excitability in a dynamical setting? In this case the size of the excitability region can be used to develop simulations of membrane potential dynamics subject to random inputs. This could be done by studying the dynamics of the size changes for *E*.

Importantly, the above results hold for both configurations of the system (3 or 1 fixed points), and for continuous models including the thermodynamical model (Herrera-Valdez, 2018), the (Hodgkin and Huxley, 1952) model, and the Fitz-Hugh model (Fitz-Hugh, 1961).

### Multi-compartment limitations and extensions

This work does not take into account the different compartments present in real neurons. Experiments and models show that sites of initiation of action potentials in neurons may move depending on factors that include the synaptic load on the dendrites (Bender and Trussell, 2012; Chen et al., 1997); but see a body of work affirming that the AP initiation site in neurons is the axon iOnenitial segment by Colbert and Johnston (1996); Foust et al. (2010); Kole and Stuart (2012) among other authors. One possible extension of the work presented here to multi-compartment models would be to calculate second time-derivatives in multi-compartment models to detect and analyse the sites where action potentials to model the location dependence of a action potential initiation.

### Excitability defined using analytical criteria across dynamical systems

A dynamical system is excitable if it has a region *R*_1_ in its phase space that contains segments of two types of trajectories, one kind that always crosses the threshold set ℐ_*θ*_, and its complement.

The excitability region *E* is thus defined analytically by calculating the level curve containing inflection points of the membrane potential with respect to time, and restricting it to *R*_1_.

Notice that this definition of excitability includes bistable systems in which one attractor is a fixed point, and one attractor is a limit cycle. However, it would not include systems in which the only attractor is a limit cycle. The reason being, that there is no way to partition the set of trajectories into two sets. That is, apart from trajectories at fixed points, all other trajectories converge to the only attractor that is an AP. Strictly speaking, such dynamical systems contain only one trajectory that converges onto a limit cycle, and will be regarded from here on as (already) “excited”.

If a dynamical system is forced further beyond having a unique limit cycle attractor to the point of having a depolarization block attractor, the system can no longer be considered as excitable, for it would not be capable of producing two types of trajectories. In such cases, all trajectories converge to a single attractor fixed point with a large *v*-value.

### The inflection curve ℐ_*θ*_ can be calculated across different continuous models

The inflection point curves shown in the sections above were calculated for a conductance-based version of a membrane potential model. However, they could be calculated for the thermodynamical model (Herrera-Valdez, 2018), or for a simple, non-biophysical model like those developed by Fitz-Hugh.

The explicit form of the equation that yields ℐ_*θ*_ is calculated using a general biophysical 2-dimensional model. Importantly, finding this threshold set ℐ_*θ*_ does not depend on whether the system is near saddle-node bifurcations or any other restrictions as previously reported (Izhikevich, 2007). It is worth noticing that a quadratic expression for *w* as a function of *v* that yields ℐ can be obtained for a particular class of 2-dimensional models (the one yielding linear K-channel activation responses in voltage-clamp, for *κ* = 0 in equation (15)).

The framework presented here unifies the theory of excitable dynamical systems for continuous models by providing an analytical description of excitability, and, *en passage* provides a tool to compare excitability across different dynamical systems defined on the same domain.

## Acknowledgments

This work was funded by the DGAPA-UNAM grant PAPIIT-IN228820. The author would like to thank Hector Chaparro-Reza, Vania Victoria Villegas-Martinez, and Lionardo Truqui for checking some of the calculations and for discussing the first ideas behind this article.

## Footnotes

1 Assume that 𝒟 is a metric space. A continuous dynamical system can be thought of as as function *φ* : ℝ × 𝒟 → 𝒟 such that (i) *φ*(0, *x*_0_) = *x*_0_ for all *x*_0_ ∈ 𝒟 (initial condition property), and (ii) *φ*(*s* + *t, x*_0_) = *φ*(*s, φ*(*t, x*_0_) (group property). If an ordinary, autonomous differential equation defined on 𝒟 has solutions for all initial conditions in, 𝒟 then the mapping describing all such solutions it can be thought of as an autonomous dynamical system (Kloeden and Rasmussen, 2011).

2 More precisely, the voltage-dependent change in the charge at the water membrane interface (in pF), usually regarded as the membrane capacitance, (Cole and Hodgkin, 1939; Herrera-Valdez, 2020)

3 The thermodynamical model for transmembrane transport (TMTT) is a unifying biophysical framework for transmembrane molecular transport derived from non-equilibrium thermodynamics that allows modelling of transmembrane flux of molecules mediated by pumps and channels with a common mathematical formulation. Conductance-based models for current can be obtained from TM expressions as approximations around the reversal potential truncated to first-order.

